# Chiron: Translating nanopore raw signal directly into nucleotide sequence using deep learning

**DOI:** 10.1101/179531

**Authors:** Haotian Teng, Minh Duc Cao, Michael B. Hall, Tania Duarte, Sheng Wang, Lachlan J.M. Coin

## Abstract

Sequencing by translocating DNA fragments through an array of nanopores is a rapidly maturing technology which offers faster and cheaper sequencing than other approaches. However, accurately deciphering the DNA sequence from the noisy and complex electrical signal is challenging. Here, we report Chiron, the first deep learning model to achieve end-to-end basecalling: directly translating the raw signal to DNA sequence without the error-prone segmentation step. Trained with only a small set of 4000 reads, we show that our model provides state-of-the-art basecalling accuracy even on previously unseen species. Chiron achieves basecalling speeds of over 2000 bases per second using desktop computer graphics processing units.

## 1 Introduction

DNA sequencing via bioengineered nanopores, recently introduced to the market by Oxford Nanopore Technologies (ONT), has profoundly changed the landscape of genomics. A key innovation of the ONT nanopore sequencing device, MinION, is that it measures the changes in electrical current across the pore as a single-stranded molecule of DNA passes through it. The signal is then used to determine the nucleotide sequence of the DNA strand^1–3^. Importantly, this signal can be obtained and analysed by the user while the sequencing is still in progress. A large number of pores can be packed into a MinlON device in the size of a stapler, making the technology extremely portable. The small size and real-time nature of the sequencing opens up new opportunities in time-critical genomics applications^4–7^ and in remote regions^8–12^.

While nanopore sequencing can be massively scaled up by designing large arrays of nanopores and allowing faster translocation of DNA fragments, one of the bottle-necks in the analysis pipeline is the translation of the raw signal into nucleotide sequence, or basecalling. Prior to the release of Chiron, basecalling of nanopore data involved two stages. Raw data series are first divided into segments corresponding to signals obtained from a k-mer (segmentation) before a model is then applied to translate segment signals into k-mers. DeepNano^13^ introduced the idea of using a bi-directional Recurrent Neural Network (RNN), that uses the basic statistics of a segment (mean signal, standard deviation and length) to predict the corresponding k-mer. The official basecallers released by ONT, nanonet and albacore (prior to version 2.0.1), also employ similar techniques. As k-mers from successive segments are expected to overlap by k-1 bases, these methods use a dynamic programming algorithm to find the most probable path, which results in the basecalled sequence data. BasecRAWller^14^ uses a pair of unidirectional RNNs; the first RNN predicts the probability of segment boundary for segmentation, while the second one translates the discrete event into base sequence. As such, basecRAWller is able to process the raw signal data in a streaming fashion.

In this article we present Chiron, which is the first deep neural network model that can translate raw electrical signal directly to nucleotide sequence. Chiron has a novel architecture which couples a convolutional neural network (CNN) with an RNN and a Connectionist Temporal Classification (CTC) decoder^15^. This enables it to model the raw signal data directly, without use of an event segmentation step. Oxford Nanopore Technologies have also developed a segmentation free base-caller, Albacore v2.0.1, which was released shortly after Chiron v0.1.

Chiron has been trained on a small data set sequenced from a viral and bacterial genome, and yet it is able to generalise to a range of genomes such as other bacteria and human. Chiron is as accurate as the ONT designed and trained Albacore v2.0.1 on bacterial and viral base-calling and outperforms all other existing methods. Moreover, unlike Albacore, Chiron allows users to train their own neural network, and it is also fully open-source, enabling development of specialised base-calling applications, such as detection of base-modifications.

## Results

### Deep neural network architecture

We have developed a deep neural network (NN) for end-to-end, segmentation-free basecalling which consists of two sets of layers: a set of convolutional layers and a set of recurrent layers (see Figure 1). The convolutional layers discriminate local patterns in the raw input signal, whereas the recurrent layers integrate these patterns into basecall probabilities. At the top of the neural network is a CTC decoder^15^ to provide the final DNA sequence according to the base probabilities. More details pertaining to the NN are provided in Methods.

**Figure 1.**
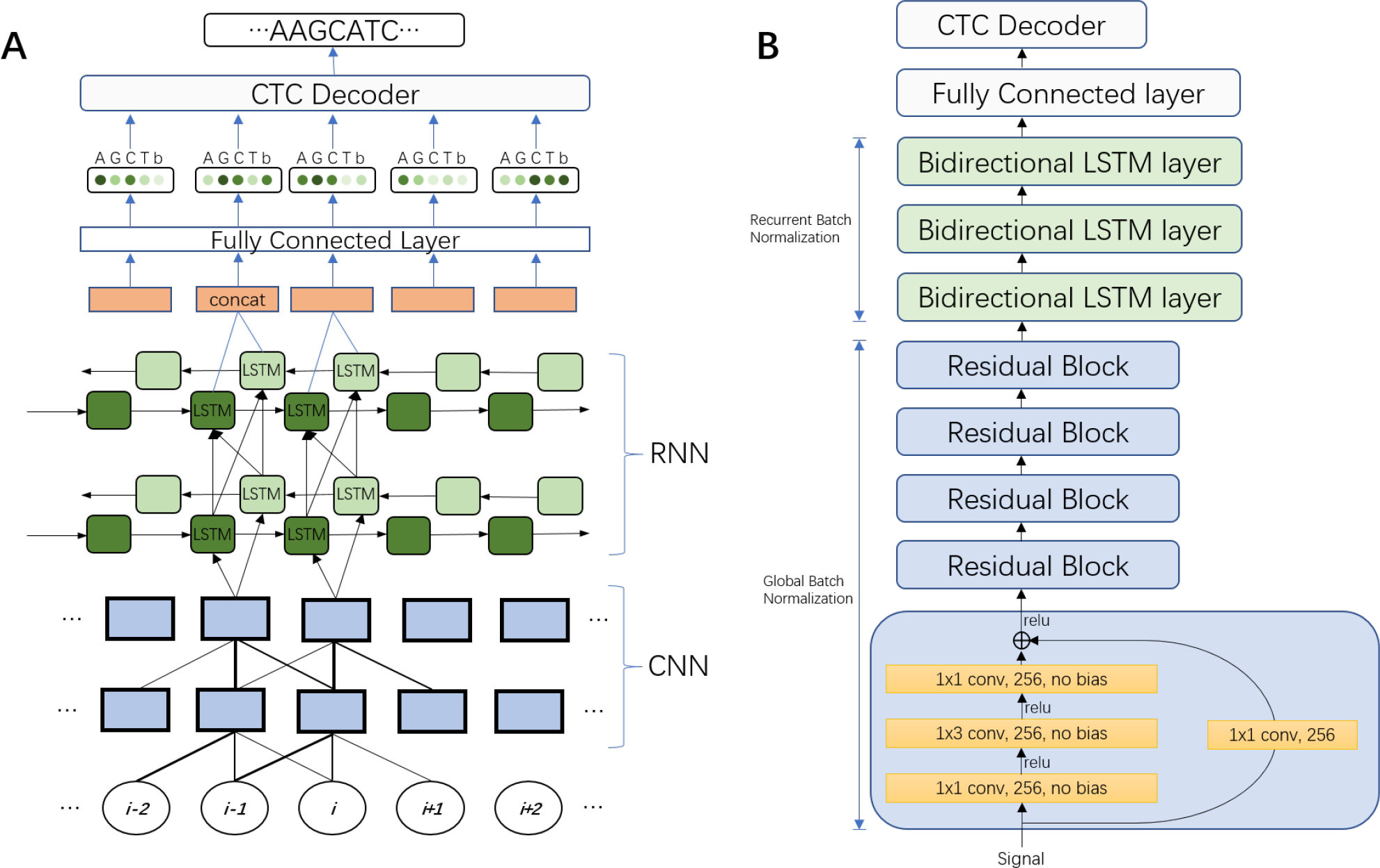
**A)** An unrolled sketch of the neural network architecture. The circles at the bottom represent the time series of raw signal input data. Local pattern information is then discriminated from this input by a CNN. The output of the CNN is then fed into a RNN to discern the long-range interaction information. A fully connected layer is used to get the base probability from the output of the RNN. These probabilities are then used by a CTC decoder to create the nucleotide sequence. The repeated component is omitted. **B)** Final architecture of the Chiron model. Variants of this architecture were explored by varying the number of convolutional layers from 3 to 10 and recurrent layers from 3 to 5. We also explored networks with only convolutional layers or recurrent layers, **1** ×3 conv, 256, no bias means a convolution operation with a 1 × 3 filter and a 256 channels output with no bias added.

**Figure 2.**
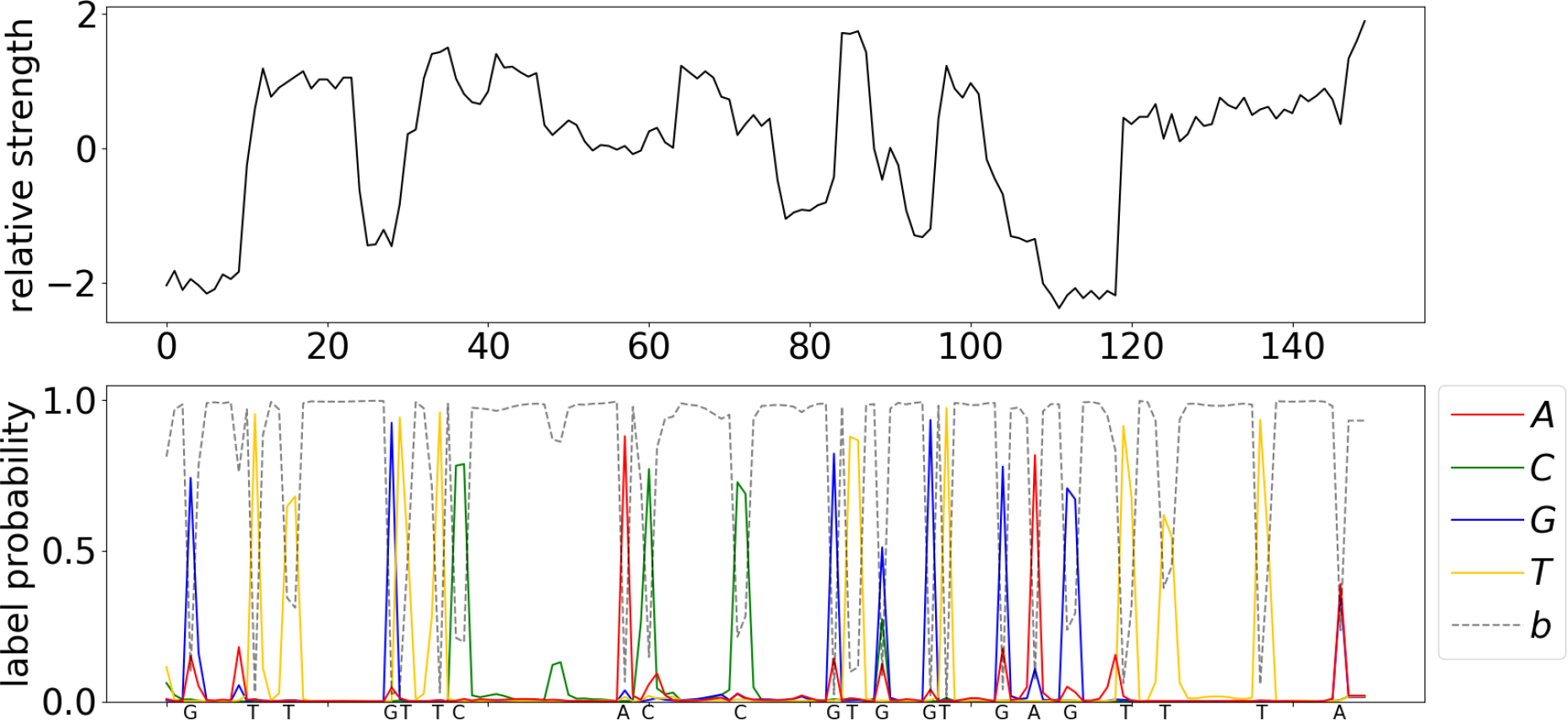
Visualization of the predicted probability of bases and the readout sequence. The upper pane is a normalised raw signal from the Minion Nanopore sequencer, normalised by subtracting the mean of the whole signal and then dividing by the standard deviation. The bottom pane shows the predicted probability of each base at each position from Chiron. The final output DNA sequence is annotated on the x-axis of the bottom plane.

Chiron presents an end-to-end basecaller, in that it predicts a complete DNA sequence from raw signal. It translates sliding windows of 300 raw signals to sequences of roughly 10-20 base pairs (which we call *slices*). These overlapping slices are stacked together to get a consensus sequence in real-time. The window is shifted by 30 raw signals, by processing this slices in parallel, the base-calling accuracy can be improved with little speed loss.

### Performance Comparison

For training and evaluating the performance of Chiron, a phage Lambda virus sample (*Escherichia virus Lambda* provided by ONT and an *Escherichia coli* (K12 MG1655) sample using 1D protocol on R9.4 flowcells are sequenced for calibrating the MinION device (See Methods). 34,383 reads were obtained for Lambda sample and 15,012 reads for *E. coli*, but only 2000 reads were randomly picked from each sample to train Chiron. It took the model 10 hours to train 3 epoch with 4,000 reads (∼ 4Mbp) on a Nvidia K80 GPU. Then Chiron is cross-validated on the remainder of the reads from two runs, and the model is further evaluated by testing its basecalling accuracy on other species. A *Mycobacterium tuberculosis* sample is sequenced and a set of human data is downloaded from chromosome 21 part 3 from the Nanopore WGS Consortium^16^, to be used in testing the generality of Chiron (see Table 7).

In order to establish the ground-truth of the data, the *E. coli* and *M. tuberculosis* samples are sequenced using Illumina technology (see Methods) and assembled, which provided a high per-base accuracy reference. The reference sequence for the Phage Lambda virus is NCBI Reference Sequence NC_001416.1 and for the human data the GRCh38 reference was used. The raw signals are labeled by identifying the raw signal segment corresponding to the nucleotide assumed to be in the pore at a given time-point (see Methods).

Table 1 presents the accuracy of the four basecalling methods, including the Metrichor basecaller (the ONT cloud service), Albacore v1.1 (ONT official local basecaller), BasecRAWller^14^ and Chiron, with a greedy decoder (Chiron) and beam search decoder(Chiron-BS), on the data. Chiron has the highest identity rate on the Lambda, *E. coli* and *M. tuberculosis* sample. Additionally, it had the lowest deletion rate, mismatch rate on Lambda, *M. tuberculosis* and *E. coli*, and the lowest insertion rate on Lambda and *E. coli*. In Human dataset where Chiron did not have the highest identity rate, it is was no more than 0.01 from the best.

**Table 1.** Results from the experimental validation and benchmarking of Chiron against three other segmentation-based Nanopore basecallers and Albacore V2(which is also segmentation-free basecaller).

| Dataset | Basecaller | Deletion Rate(%) | Insertion Rate(%) | Mismatch Rate(%) | Identity Rate(%) | Error Rate(%) |
| --- | --- | --- | --- | --- | --- | --- |
| Lambda | Metrichor | 8.93 | 2.38 | 4.57 | 86.50 | 15.88 |
|  | Albacore v1.1 | 6.35 | 3.82 | 4.69 | 88.96 | 14.86 |
|  | Albacore-v2 | **6.19** | 3.38 | **3.98** | **89.82** | 13.55 |
|  | BasecRAWller | 7.89 | 10.01 | 10.56 | 81.54 | 28.46 |
|  | Chiron | 8.20 | 2.13 | 4.03 | 87.76 | 14.36 |
|  | Chiron-BS | 6.20 | **2.13** | 4.20 | 89.60 | **12.53** |
| *E. coli* | Metrichor | 7.52 | 1.93 | 3.84 | 88.64 | 13.29 |
|  | Albacore v1.1 | 5.76 | 3.27 | 4.14 | 90.10 | 13.17 |
|  | Albacore-v2 | 5.21 | 2.99 | 3.57 | 91.22 | 11.77 |
|  | BasecRAWller | 7.16 | 10.40 | 10.30 | 82.54 | 27.86 |
|  | Chiron | 6.36 | **1.81** | **3.07** | 90.57 | 11.24 |
|  | Chiron-BS | **4.94** | 2.36 | 3.16 | **91.90** | **10.46** |
| *M. tuberculosis* | Metrichor | 7.63 | **2.40** | 4.35 | 88.02 | 14.38 |
|  | Albacore v1.1 | 6.12 | 3.57 | 4.68 | 89.19 | 14.37 |
|  | Albacore-v2 | **5.05** | 3.58 | **4.05** | **90.90** | **12.68** |
|  | BasecRAWller | 7.17 | 10.85 | 10.42 | 82.41 | 28.44 |
|  | Chiron | 7.16 | 2.50 | 4.33 | 88.51 | 13.99 |
|  | Chiron-BS | 5.84 | 3.05 | 4.50 | 89.66 | 13.39 |
| Human | Metrichor | 12.95 | **4.15** | 7.65 | 79.4 | 24.75 |
|  | Albacore v1.1 | 8.62 | 6.51 | 7.52 | 83.86 | 22.65 |
|  | Albacore-v2 | 8.71 | 6.03 | **6.05** | **85.24** | **20.79** |
|  | BasecRAWller | **8.41** | 10.28 | 10.10 | 81.49 | 28.79 |
|  | Chiron | 9.13 | 5.14 | 9.33 | 81.54 | 23.60 |
|  | Chiron-BS | 9.30 | 5.62 | 7.87 | 82.83 | 22.79 |

In addition we compared the segmentation-free ONT basecaller Albacore v2.0.1 with Chiron-BS in Table 1. Chiron-BS had a consistently lower insertion rate across all species tested, as well as a lower deletion rate on Lambda and E-coli, however it suffered a slightly higher mismatch rate on all species except E-coli. The performance is comparable to Albacore v2.0.1 on all species except for Human, however this is likely at least partially due to the fact that it has not been trained on any human DNA.

In order to assess the quality of genomes assembled from reads generated by each basecaller, we used Miniasm together with Racon to generate a de-novo genome assembly for each of the bacterial and viral genomes (see Methods). The results presented in Table 3 demonstrate that Chiron assemblies for Phage lambda and *E. coli* have approximately half as many errors as those generated from Albacore (v1 or v2) reads. For *M. tuberculosis*, Chiron has fewer errors than Albacore v1, but slightly more than Albacore v2. The identity rate and relative length for each round of polishing with Racon are shown in Figure 3.

**Figure 3.**
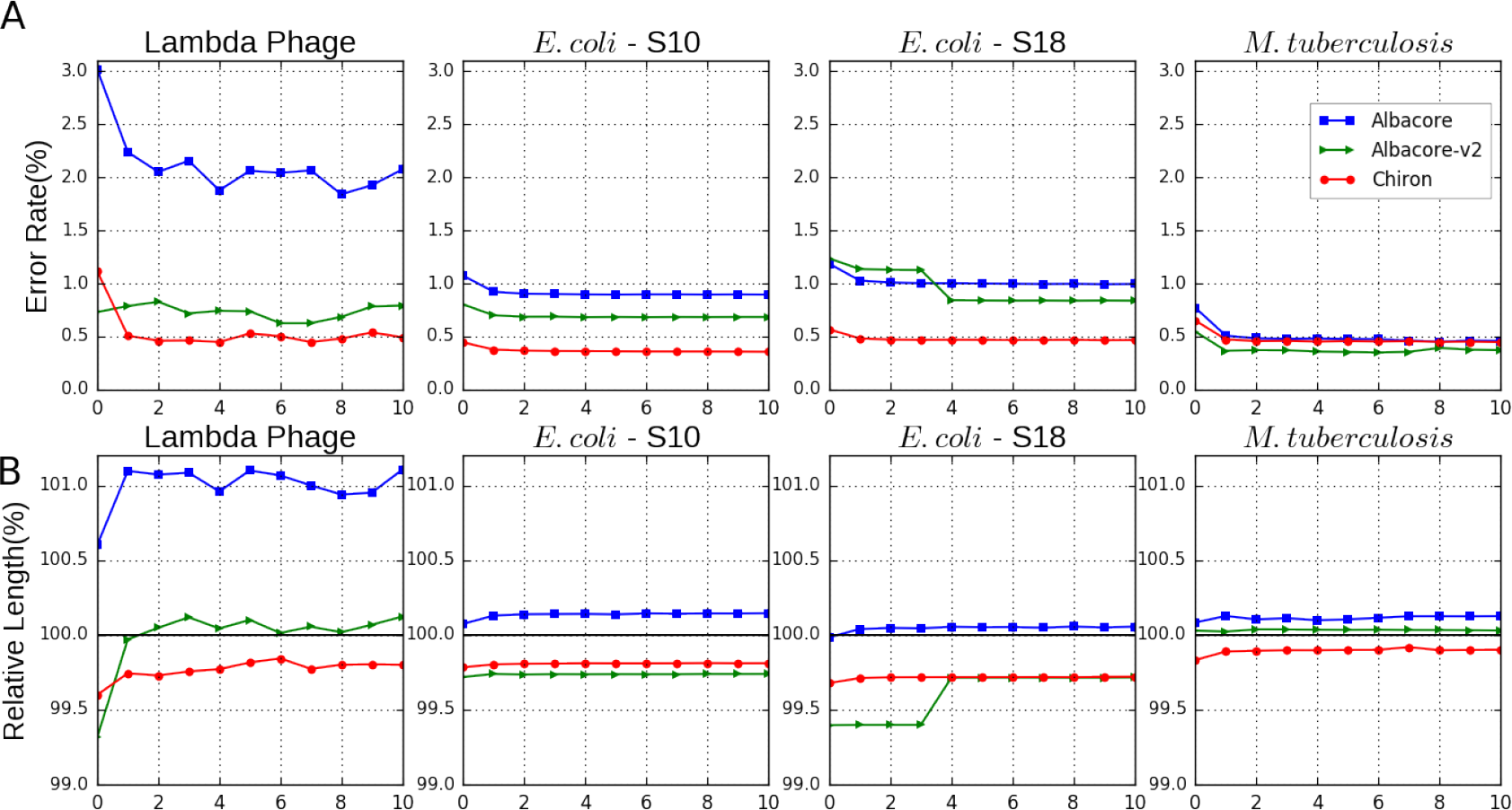
**A)** Assembly Error Rate(%) for each polishing round using Racon. Two individually sequenced *E. coli* samples are included(S10, S18). All basecallers have a similar performance on the *M. tuberculosis* dataset due to its high sequencing depth(130X). **B)** Relative assembly Length(%) after each round of polishing. *Relative length* is defined as the length of the assembly divided by the length of reference genome.

In terms of speed on a CPU processor, Chiron is slower (21bp/s, 17bp/s using a beam-search decoder with a 50 beam width) than Albacore (2975bp/s) and - to a lesser extent - BasecRAWller (81bp/s). However, when run on a Nvidia K80 GPU, a basecalling rate of 1652bp/s and 1204bp/s using a beam search decoder is achieved. (Chiron is also tested on a Nvidia GTX 1080 Ti GPU and got a rate of 2657bp/s). The GPU rate for other two local basecallers are not included, as Albacore and basecRAWller do not currently offer GPU support. Metrichor was not included in the speed benchmarking as it is not possible to gather information about CPU/GPU speed as it is a cloud basecaller.

**Table 2.** *Deletion/Insertion/Mismatch rate(%)* are defined as the number of deleted/inserted/mismatched bases divided by the number of bases in the reference genome (the lower the better), *Identity rate(%)* is defined as the number of matched bases divided by the number of bases in the reference genome for that sample (the higher the better, Identity Rate = 1 - Deletion Rate - Mismatch Rate), *Error rate(%)* is defined as the sum of deletion, insertion and mismatch rate, (the lower the better, Error Rate = Deletion Rate + Insertion Rate + Mismatch Rate). This statistic effectively summarises the basecalling accuracy of the associated model.

**Table 3.** Assembly identity rate and relative length benchmark, draft genome generated by Miniasm and is polished 10 rounds by Racon, assembly identity rates are presented in the left 4 columns while relative lengths are presented in the right 4 columns.

| Sample(coverage) | Albacore | Albacore_2 | Chiron-BS | Metrichor | Albacore | Albacore_2 | Chiron-BS | Metrichor |
| --- | --- | --- | --- | --- | --- | --- | --- | --- |
| *E. coli*-S18(27X) | 99.004 | 99.162 | 99.533 | 87.678 | 100.055 | 99.715 | 99.720 | 94.253 |
| *E. coli*-S10(40X) | 99.106 | 99.316 | 99.646 | 88.745 | 100.144 | 99.739 | 99.811 | 94.829 |
| *M. tuberculosis*(130X) | 99.541 | 99.628 | 99.554 | 84.736 | 100.126 | 100.029 | 99.900 | 90.875 |
| Lambda Phage( 790X) | 97.926 | 99.207 | 99.507 | 99.164 | 101.104 | 100.123 | 99.800 | 99.335 |

**Table 4.** *Identity rate(%)* is calculated by first shredding the assembly contigs into 10K reads pieces, and then get the mean of the identity rate of the aligned reads, *relative length(%)* is defined as the sum of the length of all the aligned pieces divided by the length of reference genome. *E. coli*-S10 and *E. coli*-S18 are reads from two independent sequencing.

**Table 5.** Base-calling rate (bp per second).

| Basecaller | CPU rate (1 core) | CPU rate (8 cores) | GPU rate |
| --- | --- | --- | --- |
| Albacorev1.1.2 | 2975 | 23800 | NA |
| BasecRAWller | 81 | 648 | NA |
| Chiron | 21 | 168 | 1652 |
| Chiron-BS | 17 | 136 | 1204 |

**Table 6.** Single core CPU rate is calculated by dividing the number of nucleotides basecalled by the total CPU time for the basecalling analysis. 8 core CPU rate is estimated by multiplying single core cpu rate by 8, based on observed 100% utility of CPU processors in multi-threaded mode on 8 cores. GPU rate calculated on a Nvidia K80 GPU. The reported rate is the average across all samples analysed. GPU rate not reported for Albacore or BasecRAWller as they have not been developed for use on GPU. Chiron is also capable of running on a GPU and its rate in this mode is included in parentheses. Albacore is not capable of running in GPU mode. Albacore V2 was found to have similar performance as albacore v1.1.2.

## Discussion

Segmenting the raw nanopore electrical signal into piece-wise constant regions corresponding to the presence of different k-mers in the pore is an appealing but error-prone approach. Segmentation algorithms determine a boundary between two segments based on a sharp change of signal values within a window. The window size is determined by the expected speed of the translocation of the DNA fragment in the pore. We noticed that the speed of DNA translocation is variable during a sequencing run, which coupled with the high level of signal-to-noise in the raw data, can result in low segmentation accuracy. As a result, the segmentation algorithm often makes conservative estimates of the window size, resulting in segments smaller than the actual signal group for k-mers. While dynamic programming can correct this by joining several segments together for a k-mer, this effects the prediction model.

All existing nanopore base callers prior to Chiron employ a segmentation step. The first nanopore basecalling algorithms^17,18^ employed a Hidden Markov Model, which maintains a table of event models for all possible k-mers. These event models were learned from a large set training data. More recent methods (DeepNano^13^, nanonet) train a deep neural network for inferring k-mers from segmented raw signal data.

A recent basecaller named BasecRAWller^14^ used an initial neural network (called a *raw* network) to output probabilities of boundaries between segments. A segmentation algorithm is then applied to segment these probabilities into discrete events. BasecRAWller then uses a second neural network (called the *fine-tune* network) to translate the segmented data into the base sequence.

Our proposed model is a departure from the above approaches in that it performs base prediction directly from raw data without segmentation. Moreover the core model is an end-to-end basecaller in the sense that it predicts the complete base sequence from raw signal. This is made possible by combining a multi-layer convolutional neural network to extract the local features of the signal, with a recurrent neural network to predict the probability of nucleotides in the current position. Finally, the complete sequence is called by a simple greedy algorithm, based on a typical CTC-style decoder^15^, reading out the nucleotide in each position with the highest probability. Thus, the model need not make any assumption of the speed of DNA fragment translocation and can avoid the errors introduced during segmentation.

To improve the basecalling speed and to minimize its memory requirements, the neural network is run on a 300-signal sliding window (equivalent to approximately 20bp), overlapping the sequences on these windows and generating a consensus sequence. Chiron has the potential to stream these input raw signal ‘slices’ into output sequence data, which will become increasingly important aspect of basecalling very long reads (100kb+), particularly if used in conjunction with the read-until capabilities of the MinION.

Our model was either the best or second-best in terms of accuracy on all of the datasets we tested in terms of read-level accuracy. This includes the human dataset, despite the fact that the model has not seen human DNA during training. Our model has only been trained on a mixture of 2,000 bacterial and 2,000 viral reads. The most accurate basecaller is the proprietary ONT Albacore basecaller. Chiron is within 1% accuracy on bacterial DNA, but only within 2% accuracy on human DNA. More extensive training on a broader spectrum of species, including human can be expected to improve the performance of our model. There are also improvements in accuracy to be gained from a better alignment of overlapping reads and consensus calling. Increasing the size of the sliding window will also improve accuracy but at the cost of increased memory and running time.

Bacterial and viral genome assemblies generated from Chiron basecalled reads all had less than 0.5% error, whereas those generated by Albacore had up to 0.8% accuracy Figure 3. This marked reduction in error rate is essential for generating accurate SNP genotypes, a pre-requisite for many applications such as outbreak tracking. These results are consistent with those reported in recent study into read and assembly level accuracy for *K. pneumoniae*^19^.

Our model is substantially more computationally expensive than Albacore and somewhat more computationally expensive than BasecRAWller. This is to be expected given the extra depth in the neural network. Our model can be run in a GPU mode, which makes computation feasible on small to medium sized datasets on a modern desktop computer. Our method can be further sped up by increasing the step size of the sliding window, although this may impact accuracy. Also there are several existing methods which can be used to accelerate NN-based basecallers such as Chiron. One such example is Quantization, which reformats 32-bit float weights as 8-bit integers by binning the weight into a 256 linear set. As neural networks are robust to noise this will likely have negligible impact of the performance. Weight Pruning is another method used to compress and accelerate NN, which prunes the weights whose absolute value is under a certain threshold and then retrains the NN^20^.

## Conclusion

We have presented a novel deep neural network approach for segmentation-free basecalling of raw nanopore signal. Our approach is the first method that can map the raw signal data directly to base sequence without segmentation. We trained our method on only 4000 reads sequenced from the simple genome lambda virus and *E. coli*, but the method is sufficiently generalised to be able to base call data from other species including human. Our method has state-of-art accuracy - outperforming the ONT cloud basecaller Metrichor as well as another 3rd-party basecaller, BasecRAWller.

## Methods

### Deep neural network architecture

Our model combines a 5-layer CNN^21^ with a 3-layer RNN and a fully connected network (FNN) in the last layer that calculates the probability for a CTC decoder to get the final output. This structure is similar to that used in speech recognition^22^. Both the CNN and RNN layers are found to be essential to the base calling as removing either will cause a dramatic drop in prediction accuracy, which is described more in the Training section.

#### Preliminaries

Let a raw signal input with *T* time-points **s** = [*s*_1_,*s*_2_,…,*s_T_*] and the corresponding DNA sequence label (with K bases) **y** = [*y*_1_,*y*_2_,…,*y_K_*] with *y_i_* ∈ {*A*,*G*,*C*,*T*} be sampled from a training dataset *χ* = {**s**^(1)^,**y**^(1)^), (**s**^(2)^,**y**^(2)^),…}. Our network directly translates the input signal time series *s* to the sequence *y* without any segmentation steps.

The input signal is normalized by subtracting the mean of the whole read and dividing by the standard deviation. 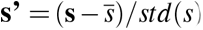.

Then the normalised signal is fed into a residual block^23^ combined with global batch normalisation^24^ in the 5 convolution layers to extract the local pattern from the signal. The stride is set as 1 to ensure the output of the CNN has the same length of the input raw signal. The residual block is illustrated in Figure 1, a convolution operation with a l× m filter, n×p stride and s output channels on a k channels input is defined as:

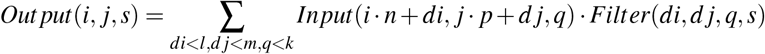

An activation operation is performed after the convolution operation. Various kinds of activation functions can be chosen, however, in this model a Rectified Linear Unit (ReLU) function is used as the activation operation which has been reported to have a good performance in CNN, defined as:

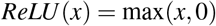

Following the convolution layers are multiple bi-directional RNN layers^25^, a LSTM cell^26^ is used as the RNN cell, with a separate batch normalisation on the inside cell state and input term^27^.

A typical batch normalisation procedure^24^ is

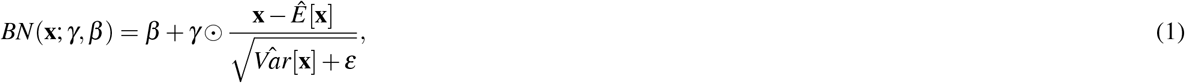

where **x** be a inactivation term.

Let 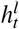 be the output of *l_th_* RNN layer at time t, the batch normalisation for a LSTM cell is

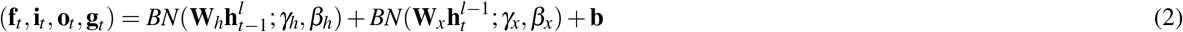

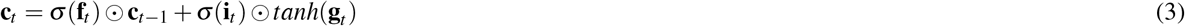

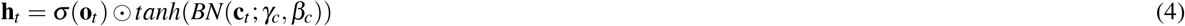

The batch normalisation is calculated separately in the recurrent term 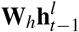 as well as the input term 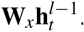 The parameters *β_h_* and *β_x_* are set to zero to avoid the redundancy with **b**. The last forward layer 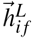 and the backward layer 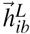 are concatenated together as an input to a fully connected layer

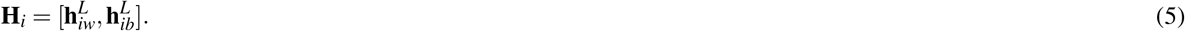

The final output is transferred through a fully connected network followed by a softmax operation

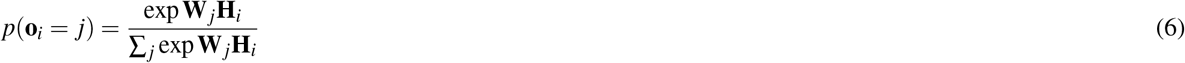

The output **o***_i_*, *i* = 1,2,…, *T* predict the symbol given the input vector **x**, *P*(*o_i_* = *j*|**x**). If the read is a DNA sequence then *j* ∈ {*A*, *G*, *C*, *T*, *b*}, where b represents a blank symbol(Figure 1). During training, the CTC loss is calculated between the output sequence **o** and label **y**^15^ and back-propogation is used update the parameters. An Adam optimizer^28^ with an initial learning rate of 0.001 is used to minimize the CTC loss.

During inference, the final sequence constructed using either a greedy decoder^15^, or a beam-search decoder^29^. The greedy decoder works by first getting the argument of maximum probability in each position of **o**, and then producing the sequence call by first removing the consecutive repeat, and then removing the blank symbols. For example, the greedy path of an output **o** is A A - - - A - - G -, here - represent the blank symbol, the consecutive repeat is removed first and lead to A - A - G -, and the blank is removed to get the final sequence AAG. The beam search decoder with beam width W, maintains a list of the W most probable sequences (after collapsing repeats and removing blanks) up to position i of **o**. To obtain this list at position i+1, it constructs the probability of all possible extensions of the W most probable at position i based on adding each symbol according to *p*(*o_i_* = *j*), and collapsing and summing up over repeated bases, or repeated blanks which are terminated by a non-blank. The greedy decoder is a special case of the beam-search decoder when the beam width is 1. It should be noted that the model can still call homopolymer repeats provided each repeated base is separated by a blank, which is typically the case.

#### Convolutional network to extract local patterns

256 channel filters are used for all five convolutional layers. In each layer, there is a residual block^23^ (Figure 1) composing with two branches. A 1×1 filter is used for reshaping in the first branch. In the second branch, a 1×1 convolution filter is followed by a rectified linear unit (RELU)^30^ activation function and a 1×3 filter with a RELU activation function as well as a 1×1 filter. All filters have the same channel number of 256. An element-wise addition is performed on the two branches followed by a RELU activation function. A global batch normalisation operation is added after every convolution operation. A large kernel size (5,7,11) and different channel numbers (128,1024) is also tested, and the above combination is found to yielded the best performance.

#### Recurrent layers for unsegmented labelling

The local pattern extracted from the CNN described above is then fed to a 3-layer RNN (Figure 1). Under the current ONT sequencing settings, the DNA fragments translocate through the pore with a speed of roughly 250 or 450 bases per second, depending on the sequencing chemistry used, while the sampling rate is 4000 samples per second. Because the sampling rate is higher than the translocation rate, each nucleotide usually stays in the current position for about 5 to 15 samplings, on average. Furthermore, as a number of nearby nucleotides also influence the current, 40 to 100 samples (based on a 4- or 5-mer assumption) could contain information about a particular nucleotide. A 3-layer bidirectional RNN is used for extracting this long range information. LSTM (Long Short Term Memory) cells^31,32^ with 200 hidden units are used in every layer and a fully connected neural network (FNN) is used to translate the output from the last RNN layer into a prediction. The output of the FNN is then fed into a CTC decoder to obtain the predicted nucleotide sequence for the given raw signals.

#### Improving basecalling performance

To achieve a better accuracy and less memory allocation, a sliding window is applied (default of 300 raw signals), with a pre-set sliding step size (default of 10% of window size), to the long raw signal. This gives a group of short reads with uniform length (window length) that overlap the original long read. Then basecalling is run in parallel on these short reads, and reassemble the whole DNA sequence by finding the maximum overlap between two adjacent short reads, and read out the consensus sequence. Note here the reassembly is very easy because the order of the short reads is known. This procedure improves the accuracy of the basecalling and also enables parallel processing on one read.

### Data preparation

#### Sequencing

The library preparations of the *E. coli* and *M. tuberculosis* samples were done using the *1D gDNA selecting for long reads using SQK-LSK108* (March 2017 version) protocol with the following modifications. Increase the incubation time to 20 minutes in each end-repair and ligation step; use 0.7x Agencourt^R^ AMPure^R^ XP beads (Beckman Coulter) immediately after the end-repair step and incubation of the eluted beads for 10 minutes; and use elution buffer (ELB) warmed up at 50°C with the incubation of the eluted bead at the same temperature. For the Lambda sample, the *1D Lambda Control Experiment for MinION^TM^ device using SQK-LSK108* (January 2017 version) protocol was followed with some changes: sheared the sample at 4000rpm (2×1 minutes); 30 minutes of incubation in each end-repair step and 20 minutes for adaptor ligation and elution of the library with 17*μ*L of ELB. All samples were sequenced on new FLO-MIN106, version R9.4, flow cells with over 1100 active single pores and the phage was sequenced in a MinION Mk1 (232ng in 6h run) while the bacteria samples were sequenced in a MinION Mk1B (1*μ*g *E. coli* and 595ng *M. tuberculosis* in 22h and 44h runs, respectively). The *E. coli* sample was run on the MinKNOW version 1.4.3 and the other samples in earlier versions of the software. The *E. coli* sample was also sequenced on Illumina MiSeq using paired-end 300×2 to 100-fold coverage. An assembly of the *E. coli* genome was constructed by running Spades^33^ on the MiSeq sequencing data of the sample. The genome sequence of the Phage Lambda is NCBI Reference Sequence: NC_001416.1.

#### Labelling of raw signal

Metrichor, the basecaller provided by ONT which runs as a cloud service, is used to basecall the MinION sequencing data first. Then Nanoraw^34^ is used for labelling the data. Briefly, the basecalled sequence data is aligned back to the genome of the sample, and from the alignment the errors introduced by Metrichor are corrected to avoid the bias from Metrichor being learned into Chiron, and the corrected data is mapped back to the raw data. The resulting labelling consists of the raw signal data, as well as the boundaries of raw signals when the DNA fragment translocates to a new base. We use the base-level segmentation of the raw data to obtain matched pairs of signal segment (of lengths 200, 400 and 1000) together with the corresponding DNA base sequence. From this point onwards, the exact matching of the signal to each base within a segment is disregarded.

#### Training dataset

A data set using 2,000 reads from *E. coli* and 2,000 reads from Phage Lambda is created for training Chiron. In every start of the training epoch, the dataset is shuffled first and then fed into the model by batch. Training on this mixture dataset gave the model better performance both on generality and accuracy on not only the *E. coli* and Phage Lambda but also on *M. tuberculosis* and Human data.

### Training

The labelling from Metrichor described previously in Equation 1 is used to train Chiron, although the neural network architecture is translation invariant and not restricted by the sequence length, a uniform length of sequences is suited for batch feeding, thus can accelerate the training process. From this view, the original reads were cut into short segments with a uniform length of 200, 400 and 1000, and trained on these batches in alternation. Several different architectures of the neural network were tested, (see Table 8) with the CNN-RNN network architecture having the best accuracy compared to a CNN- or RNN-only network. Also using more layers seems to increase the performance of the model, however, the time consumed for training and basecalling is also increased. In the final structure, a NN with 5 convolution layers and 3 recurrent layers is adopted, as adding layers above this structure gave negligible performance improvement but required more calculation and also increased the risk of overfitting.

**Table 7.** Details about the number of reads and their median read length for data that was used in evaluation of the various basecallers.

| Sample | No. reads | Median read length (bp) |
| --- | --- | --- |
| Phage Lambda | 34,383 | 5720 |
| *E. coli* | 15,012 | 5,836 |
| *M. tuberculosis* | 147,594 | 3,423 |
| Human | 10,000 | 6,154 |

**Table 8.** Comparison of normalised edit distance with different neural network architectures. The normalised edit distance is the edit distance between predicted reads and labelled reads and normalised by segments length.

| Architecture | normalised edit distance |
| --- | --- |
| 3 Convolutional Layers | $0.4007 \pm 0.0277$ |
| 5 Convolutional Layers | $0.3903 \pm 0.0230$ |
| 10 Convolutional Layers | $0.3874 \pm 0.0186$ |
| 3 Bidirectional Recurrent Layers | $0.2987 \pm 0.0221$ |
| 5 Bidirectional Recurrent Layers | $0.2930 \pm 0.0215$ |
| 3 Convolutional Layers + 3 Bidirectional Recurrent Layers | $0.2011 \pm 0.0252$ |
| 5 Convolutional Layers + 5 Bidirectional Recurrent Layers | $0.2001 \pm 0.0177$ |

### Parameters for basecalling

All basecallers were invoked on the same set of reads for each sample. When using Chiron to basecall, the raw signal was firstly sliced by a 300 length window, the window is slided by 30, and then these sliced segments are fed into the basecaller with a batch size equal to 1100, and then the output short reads are simply assembled by a pair-wise alignment between neighbouring reads, and the consensus sequence is output from this alignment. All basecalling with Albacore (version 1.1.1 and version 2.0.1) and BasecRAWller^14^ (version 0.1) was done with default parameters. For the configuration setting in Albacore, `r94_450bps_linear.efg` was used for all samples, as this matches the flowcell and kit used for each sample. The data is basecalled on Metrichor on Jun 3rd 2017(Lambda), May 18th 2017(E. *coli)*, Jun 4th 2017(M. *tuberculosis)*, and June 20th 2017(NA12878-Human).

### Quality score

The quality score is calculated by the following algorithm: 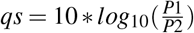 where P1 is the probability of most probable base in current position, and P2 is the probability of the second probable base in current position.

### Comparison of raw read accuracy

To assess the performance of each program, the resulting FASTA/FASTQ file from basecalling was aligned to the reference genome using graphmap^35^ with the default parameters. The resulting BAM file is then assessed by the `japsa` error analysis tool (`jsa.hts.errorAnalysis`) which looks at the deletion, insertion, and mismatch rates, the number of unaligned and aligned reads, and the identification rate compared to the reference genome. The identity rate is calculated as 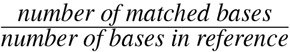 and is the marker used here for basecalling accuracy.

### Assembly Identity Rate Comparison

We assessed the quality of assemblies generated from reads produced by different base-callers. For each base-caller, a de-novo assembly is generated by the use of only Nanopore reads for the *M. tuberculosis E. coli* and Lambda Phage genomes. We use Minimap2^36^ and Miniasm^37^ to generate a draft genome, then Racon^38^ is used to polish on the draft genome for 10 rounds.

### Data availability

The *M. tuberculosis* sequencing data have been deposited Genbank under project number PRJNA386696. The Human nanopore data were downloaded from https://github.com/nanopore-wgs-eonsortium/NA12878. The *E. coli* data are in the process of being deposited to Genbank.

Program and code are available at https://github.com/haotianteng/chiron pypi package index 0.3 at https://pypi.python.org/pypi/chiron. Chiron is registered in SciCrunch with RRID:SCR_015950.

## Authors’ contributions

MH, MDC and LC conceived the study and designed the experimental framework. HT designed and implemented the Chiron algorithm. MDC, LC and TD designed and performed the MinlON sequencing. HT and MDC labelled the training data. HT and MH ran the performance comparison. HT and MDC wrote the initial draft. HT, MH and LC refined the manuscript. All authors contributed to editing the final manuscript.

## Acknowledgements

LC is supported by an NHMRC career development fellowship (FT110100972). The research is supported by an ARC research grant (DP170102626). MH is supported by a Westpac Future Leaders Scholarship (2016) awarded by the Westpac Bicentennial Foundation. We thank Jianhua Guo for contributing the DNA for the *E. coli* sample. We thank Arnold Bainomugisa for extracting DNA for the *M. tuberculosis* sample. We thank Sheng Wang and Han Qiao for the helpful discussion. We would like to thank Jain et al.^16^ for the open human nanopore dataset

